# Identification of dialects and individuals of globally threatened Yellow Cardinals using neural networks

**DOI:** 10.1101/2023.06.07.544140

**Authors:** Hernan Bocaccio, Marisol Domínguez, Bettina Mahler, Juan Carlos Reboreda, Gabriel Mindlin

## Abstract

Audio-based analysis of bird songs has proven to be a valuable practice for the growth of knowledge in the fields of ethology and ecology. In recent years, machine learning techniques applied to audio field recordings of bird calls have yielded successful results in studying population distributions and identification of individuals for their monitoring in a variety of bird species. This offers promising possibilities in the study of social behavior, biodiversity, and conservation strategies for birds. In this work, we trained deep learning models, directly from the sonograms of audio field recordings, to investigate the statistical properties of vocalizations in the endangered Yellow Cardinal. To our knowledge, this is the first time that this approach has been applied to this threatened species. Our results indicate the presence of vocal signatures that reflect similarities in songs of individuals that inhabit the same region, determining dialects, but which also show differences between individuals. These differences can be exploited by a deep learning classifier to discriminate the bird identities through their songs. Models trained with data labeled by regions showed a good performance in the recognition of dialects with a mean accuracy of (0.84 +/-0.04) significantly higher than accuracy obtained by chance. Precision and recall values also reflected the classifier’s ability to find alike vocal patterns in the songs of neighboring individuals. Models trained with data labeled by individuals showed an accuracy of (0.63 +/-0.03) significantly higher than that obtained by chance. However, the individual discrimination model showed greater confusion with neighboring individuals. This reflects a hierarchical structure in the characteristics of the Yellow Cardinal’s vocalization, where the intra-individual variability is lower than the inter-individual variability, but it is even lower than the variability obtained when individuals inhabit different regions, providing evidence of the existence of dialects. This reinforces the results of previous works but also offers an automated method for characterizing cultural units within the species. Along with genetic data, this method could help better define management units, thereby benefiting the success of reintroduction programmes of recovered individuals of Yellow Cardinal. Moreover, the novelty of individual discrimination using neural networks for the Yellow Cardinal, which has limited data availability, shows promise for non-invasive acoustic monitoring strategies with potentially relevant implications for its conservation.

## 1. Introduction

The Yellow Cardinal *Gubernatrix cristata* is currently listed as an endangered species according to the International Union for Conservation of Nature (IUCN) Red List (BirdLife International, 2018). This species of passerine bird, from the Thraupidae family (Barker et al., 2013), is endemic to the southern regions of South America (Ridgely and Tudor, 1989), including the south of Brazil (Beier and Fontana, 2019; Beier et al., 2017; Bülau et al., 2021), Uruguay (Azpiroz et al., 2012; Domínguez et al., 2017), and central Argentina, where the largest populations were found (Domínguez et al., 2020, 2017). The global population is estimated to be between 1000 and 2000 mature individuals (BirdLife International, 2018), distributed in small and fragmented populations (Collar et al., 1997). Current studies show that the Yellow Cardinal has suffered a drastic and persistent population decline throughout its range (Domínguez et al., 2020; Reales et al., 2019). This is attributed to the loss of natural habitat and to the capture of birds for illegal trade and mascotism (Domínguez et al., 2020; Pessino and Tittarelli, 2006; Segura et al., 2019). Because bird trapping is focused on the extraction of males for use as cage birds (Pessino and Tittarelli, 2006), probably due to the richer vocalizations of males and the characteristics of the male’s plumage (Chebez, 1994; Collar et al., 1992; Ridgely and Tudor, 1989), a lack of males is assumed in Yellow Cardinal populations (Segura et al., 2019). This lack of males could be cause of the hybridization with the Common Diuca in some regions (Bertonatti and López Guerra, 1997; Domínguez et al., 2016), as well as observed breeding anomalies like polygyny (Segura et al., 2019) that hinder reproductive success. Additionally, another factor affecting the reproduction of the species is the impact of both Shiny Cowbird and botfly parasitism, resulting in brood reduction and often in nest abandonment (Azpiroz, 2015; Domínguez et al., 2015), which has been shown to even lead to zero recruitment in a breeding season for this species (Atencio et al., 2022). Therefore, the status of this threatened species becomes more worrying every day, and more and diverse strategies should be considered in pursuit of its conservation. In this way, the exploitation of computational techniques, such as audio-based analysis of bird songs from field recordings, could be very helpful for effective non-invasive monitoring and conservation planning.

In the last decade, the analyses of animal vocalizations from audio recordings, improved by the application of computational methods from data science and deep learning (LeCun et al., 2015), have become prominent techniques for addressing multiple animal behavior questions. The most visible precedent is the Bird-CLEF contest (Goëau et al., 2014) in which several advances in the recognition of bird species were achieved through their songs extracted from audio data collected by the crescent Xeno-Canto initiative (Vellinga and Planqué, 2015). Publications related to these competitions have shown successful approaches for that task, allowing progress in the study of biodiversity (Goëau et al., 2016), particularly since convolutional neural networks started to be used as models trained from images representing the temporal evolution of acoustic features (Piczak, 2016; Sprengel et al., 2016; Tóth and Czeba, 2016). This approach has become the main one for bird species recognition and, although large amounts of data are often used, in many cases data augmentation techniques analogous to those used for visual tasks are applied (Kahl et al., 2017). Recently, deep learning techniques started to be applied for the identification of individual birds through their songs (Bedoya and Molles, 2021; Bistel et al., 2022a,b; Martin et al., 2022; Tubaro and Mindlin, 2019). The vocal recognition to track individuals within a population requires that bird call features show lower withinthan between-individual variation and that this variation is stable over the course of an individual’s life (Budka et al., 2015). This could be useful for monitoring the social behavior of a small population, which may be particularly relevant in the case of threatened species, where songs play a fundamental role in a variety of social interactions, from territorial defense to partner selection (Bistel et al., 2022b). More generally, the recognition capacity of these models is conditioned to the statistical properties of data in such a way that among class variability must be higher than within class variability, and a classifier has to be able to capture and exploit these differences.

Since imitative vocal learning allows for the generation and rapid transmission of new patterns of vocal structure (Slabbekoorn and Smith, 2002; Slater, 1989), it is plausible to find differences in songs between populations of the same species (Catchpole and Slater, 2008) that inhabit different geographical regions. This results in the presence of local songs, often described as dialects (Lemon, 1975), which occur when a group of conspecific males share vocal characteristics that differ slightly from those of other groups (Baker and Cunningham, 1985). It is hypothesized that these geographical variations could be caused by the interaction between learning and isolation mechanisms and influenced by the effects of cultural drift, genetic drift, cultural selection, natural selection, and sexual selection (Catchpole and Slater, 2008; Podos et al., 2004; Podos and Warren, 2007). Due to the fragmentation of the environment and the isolation of the populations of Yellow Cardinals (Domínguez et al., 2020), the presence of song dialects is expected. In fact, previous studies have reported evidence of dialects in the songs of the Yellow Cardinal (Domínguez et al., 2016), and males of the Yellow Cardinal species can recognize these dialects since they respond more strongly to local songs than to foreign ones in playback experiments (Fracas et al., 2023). This could potentially affect reproductive behavior in the Yellow Cardinal and highlights the need to understand the properties of cultural units, which is a sensitive issue for developing better strategies for the reintroduction of individuals of threatened species.

Here we used deep learning models to build sonogram-based classifiers for the study of audio field recordings of Yellow Cardinal vocalizations in individuals-labeled data from four geographical regions. We first trained a model to detect from which region a vocalization is, by assigning a unique label to all audios from individual birds that inhabit the same region. Therefore, this classifier is oriented to the study of the conformation of dialects in Yellow Cardinal songs, to reinforce the results obtained in previous works (Domínguez et al., 2016; Fracas et al., 2023), but from a deep learning techniques perspective. Then, we trained a model to discriminate individuals, from audio labeled with bird identities. The aim of this work is to use deep learning techniques that allow the identification of dialects and individuals, and contribute to a non-invasive monitoring and conservation planning of the species.

## 2. Material and Methods

### 2.1. Data

Audio data reported in a previous work was used (Domínguez et al., 2016), acquired during four breeding seasons (September-January 2011-14) in four different populations covering most of the known distribution of the Yellow Cardinal in Argentina and Uruguay: Corrientes (Co), San Luis (SL), Rio Negro (RN) and Uruguay (U) (Figure 1). Differences among these regions were found in terms of genetics (Domínguez et al., 2017) and in song dialects (Domínguez et al., 2016). Recordings were obtained using a Zoom ZOH4NK H4n Handy Recorder in close proximity to singing birds (5-10 m), as stereo or mono audios with 44100 Hz or 96000 Hz sampling rate and PCM 16-bits resolution. Birds had been ringed previously, allowing the association of each audio with an individual bird. Poor quality recordings were discarded, as well as those songs of individuals with less than three songs. The dataset finally used in this work included three to fourteen (5.08 +/-2.67) high-quality songs per individual of 25 males (8 from Co, 6 from RN, 3 from SL, and 8 from U), formed by 127 songs (54 from Co, 27 from RN, 13 from SL, 33 from U) with durations of between 2 and 11.3 (4.50 +/-1.39) seconds.

**Figure 1:**
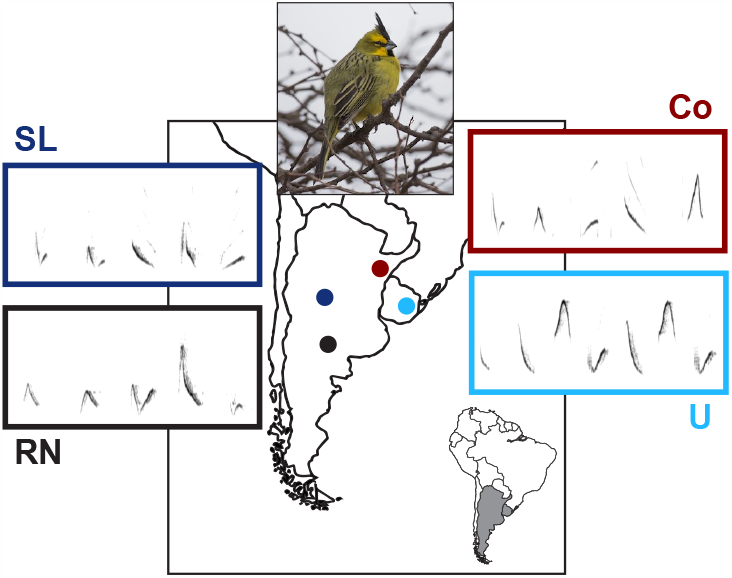
Data. Map of South America showing the four regions of Yellow Cardinals recordings included in this study, along with examples of songs spectrograms of individuals representing each population. The color code used for the population regions is as follows: Corrientes (Co) in red, San Luis (SL) in blue, Rio Negro (RN) in black, and Uruguay (U) in sky blue. Photo from Chris Wood (Macaulay Library at the Cornell Lab of Ornithology; ML31529631).

### 2.2. Data pre-processing

The full set of audio files was first converted to mono and downsampled to 22050 Hz. A 12th-order Butterworth FIR type band-pass filter was applied, with cut-off frequencies between 1 and 10 kHz. A noise reduction filter is then applied by using spectral gating as a form of noise gate which uses a threshold for each frequency band and suppresses the corresponding spectral component of noise below this threshold. For this purpose, stationary estimation of noise thresholds was made with parameters selected for a signal and noise threshold placed in 1.5 standard deviations above the mean and then the suppression with a noise reduction proportion of 95%. Thereupon, the audios were divided into 3-second chunks with a 2-second overlap, padding with zeros when chunks did not last 3 seconds, but discarding chunks that needed more than 1 second of padding. Then, spectrograms for each chunk were calculated by applying the Short-Time Fourier Transform (STFT), using a Hann window function with a window size of 256 time samples, an overlap of 64 time samples (25%), and a hop-size of 192 time samples (75%). The length of the FFT used was of 256 frequency samples, resulting in a spectrogram shape of (129, 344) matrix elements. Clipping operation was applied to spectrograms, limiting values to the range between 3 times the median (values below was set to zero) and the maximum. Spectrograms were flipped for convention. The logarithm values were computed for each element of the spectrogram matrix. Finally, spectrograms were rescaled between 0 and 1. Preprocessing steps are shown in Figure 2.a., and were made in Python by using scipy and noisereduce (Sainburg, 2019) packages.

**Figure 2:**
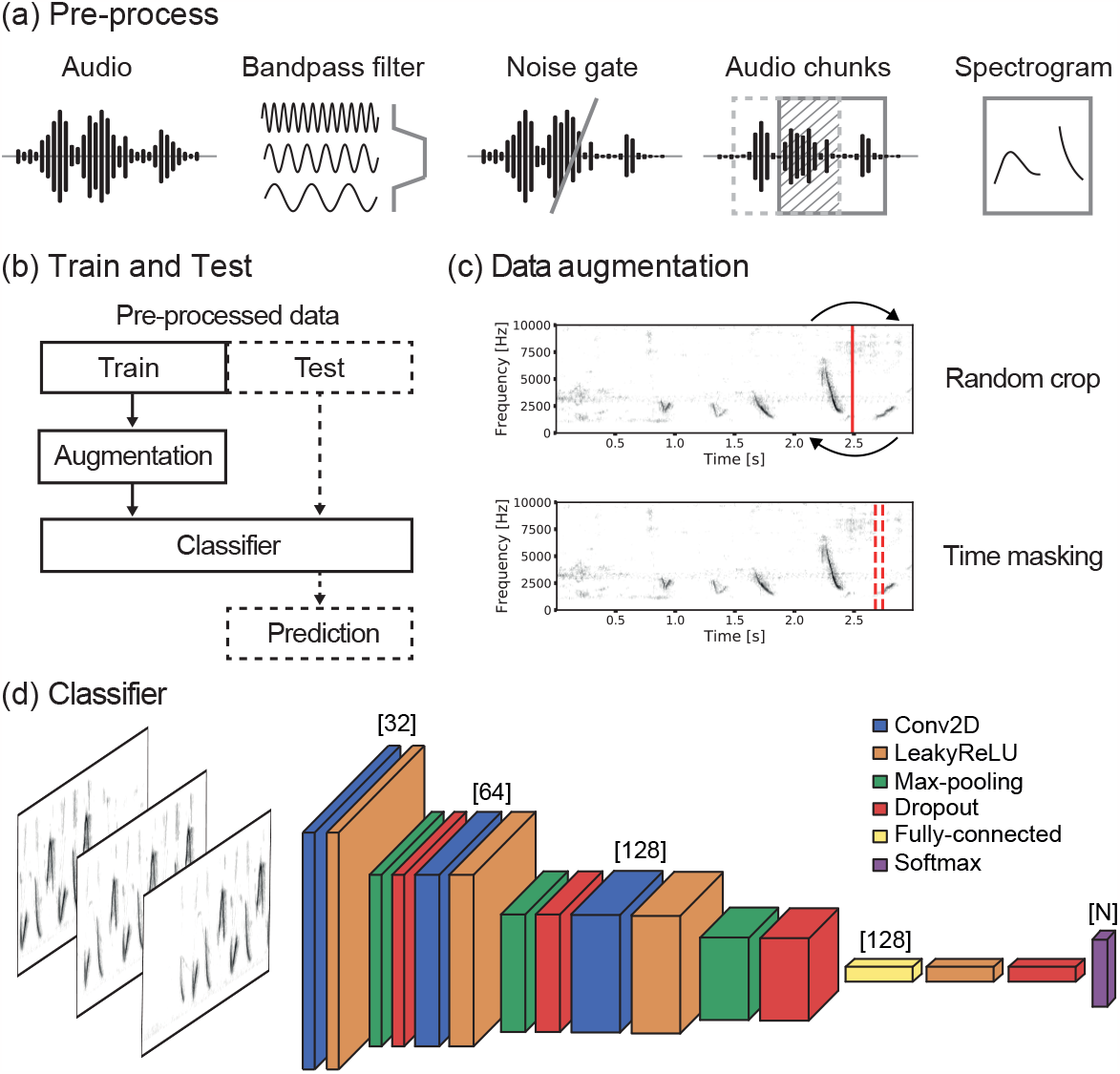
Materials and Methods. (a) Pre-process steps of audio data and construction of spectrograms used. (b) Training and test data splitting. Data augmentation was applied only to the training set. Predictions were made only on the test set. Data augmentation techniques used. Random crop was made by randomly selecting a time mark and then swapping the spectrogram pieces that were before and after the mark. Time masking was made by randomly selecting a range of times and then reducing the intensity of the spectrogram by a random factor between 0 and 1 in that range. (d) Classifier trained using spectrogram chunks as inputs, consisting in a convolutional neural network with N outputs corresponding to the number of classes.

**Figure 3:**
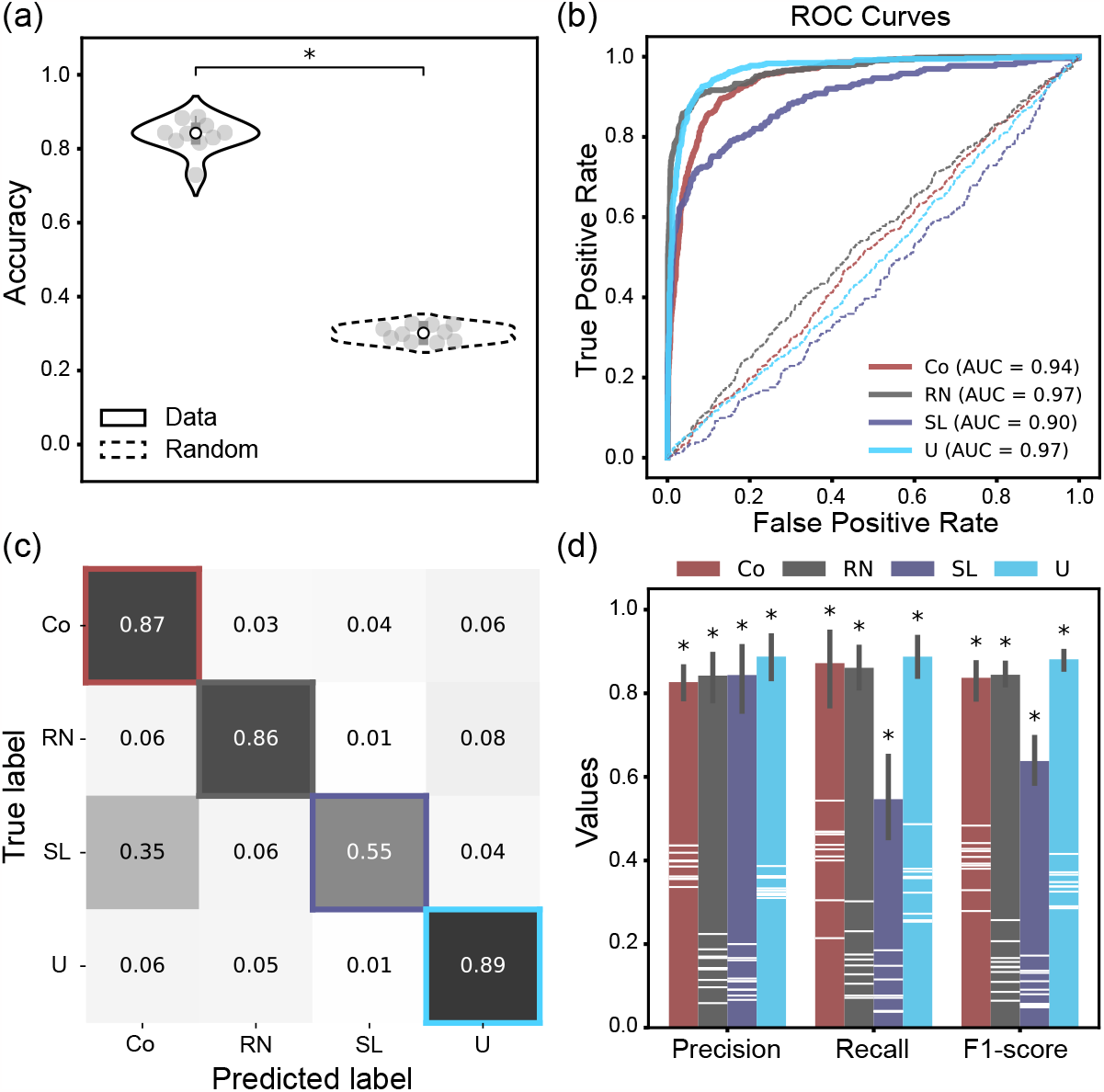
Results of dialects classifiers. Population color code used for plotting (Co: red; RN: black; SL: blue; U: sky blue) for the regions of Corrientes (Co), San Luis (SL), Rio Negro (RN) and Uruguay (U). (a) Violin plot of models accuracies obtained for test data (solid line) and random labeled test data (dashed line), reflecting significant higher values for the model (*; p-value=2.82e-11) by comparing with a paired sample t-test. (b) ROC curves of the classifier for each class against the others (solid lines) with their respective ROC AUC scores (Co: 0.94; RN: 0.97; SL: 0.90; U: 0.97), and those obtained by chance (dashed lines), showing good results of the classifier to discriminate songs among regions. (c) Confusion matrix of the classifier. The model showed good performances in prediction for most classes, with the exception of San Luis (SL) class, for which showed not bad performance but with some undesirable confusions mostly with Corrientes (Co) class. Precision, Recall, and F1-score values obtained for the different regions. All performances values (bars plot with error) were significantly higher than those obtained by chance (white lines inside the bars) according to Wilcoxon signed-rank tests (*; p-value < 0.05).

### 2.3. Train and Test

After pre-process acoustic data from the 25 male individuals, the full set of spectrogram chunks was randomly divided into training and test sets. The data division was done in such a way as to ensure that all the spectrograms associated with a song belonged to the same set and that the test set contained at least 50% of the songs of each individual. Then, the test set was formed by a random selection of half (half plus one) of songs for each individual and the chunks belonging to these songs, when the total number of songs was even (odd). Before training, data augmentation operations were applied to the selected training set. The augmented pool of spectrogram chunks was split to reserve 25% for validation, to evaluate and control model overfitting, and the rest for actually training the model. Once the model was trained using as inputs the spectrograms resulting from the augmented data, the classifier was applied to the independent test set, which was not submitted to data augmentation, for obtaining the model predictions. The scheme of the workflow used could be found in Figure 2.b.

### 2.4. Data augmentation

We increased training data to improve the model’s capacity. Common data augmentation operations implemented to visual tasks would create artificial sonograms that, while could enhance the performance of the trained models, do not reflect biologically reliable data (e.g. rotations). We constrained the augmentation techniques to which ones we considered could be plausible to find in field recordings either by audio artifacts or by intrinsic biological variability. We first used random crop in time technique, by which a randomly selected time defines a time mark, and then a swap is performed in the order of the subseries positioned before and after the crop mark. To avoid undesired artifactual syllables in bird songs, obtained from cropping of syllable fragments that could have been previously cut in the step of chunk splitting or due to the recording properties, we selected random time marks sufficiently separated from chunk edges. We then applied a time masking, reducing intensity of the spectrogram by a random factor between 0 and 1 for a range of times, starting from a randomly selected time position, and with a random range width defined to that intensity reduction could only affect a section of a syllable and not multiple syllables. We generated 20 augmented instances from each spectrogram chunk, by applying each of these augmentation steps with 90% of probability. Hence, we increased data by using spectrogram transformations compatible with audio sampling properties, as shown in Figure 2.c.

### 2.5. Convolutional neural network

The classifier consists of a convolutional neural network (CNN) implemented using Keras package in Python (Chollet, 2021), with 3 convolutional layers composed of 32, 64, and 128 filters, respectively. For all convolutions we use a 3x3 kernel, and we define the step size with which the filter moves across the image (“stride”) as 1 in both directions and the “padding” in such a way that the activation maps have the same dimensions as the input for this kernel size. Subsequently to each convolutional layer we add a layer where a LeakyReLU activation function is applied, which is similar to the ReLU activation function but instead of a flat slope it has a small slope for negative values that we configure as 0.1, followed by a layer of sampling reduction from “max-pooling”. The max-pooling operation finds the maximum value between a sample window and passes this value as a summary of characteristics over that area. As a result, the size of the data is reduced by a factor equal to the size of the sample window on which it operates, which in this case was 2x2, so the activation maps reduce their dimensions by half.

In cases where the dimension of the activation map is odd and the half does not give an integer value, a row or column of pixels of value 0 (“padding”) is added to make the dimension even. In this way, in those cases, the dimension of the map remains (d+1)/2 instead of d/2. Since input images have a shape of (129, 344) and the network architecture consists of 3 pooling layers, “padding” was used to reduce the sampling in all layers only for the first dimension. To reduce overfitting, we also added a “dropout” layer at the output of each sample reduction layer, where a fraction of random neurons (0.5, 0.5, and 0.4, respectively) were left out of training. After these 3 convolutional blocks, we add a block made up of a layer of 128 densely connected neurons (“fully-connected”) with a LeakyReLU activation function with a negative slope of 0.1 and a “dropout” of 0.3. The network finishes with a layer of N neurons (where N is the number of classes) with activation function “softmax” (a generalization of the logistic function), which allows the multi-class classification of the data. The output of each of these neurons is the probability of belonging to one of the N classes, normalized so that the sum of the outputs of an instance equals 1. The architecture of the model is shown in Figure 2.d. The model parameters were optimized by applying backward propagation to update the adjusted values during a process of 10 training epochs, starting from a predetermined initialization from the Glorot method with uniform distributions (Glorot and Bengio, 2010). We use the categorical cross entropy as a loss function, minimized during training using Adam’s algorithm (Kingma and Ba, 2014), which is a method to apply stochastic gradient descent in an optimized way. The loss function computation for the following update of model weights was made by grouping training inputs using the mini-batch method with a batch size of 32 instances. We checked that both the amount of well classified data and the cost function calculated on the training and validation groups show that models were not overfitting the data.

### 2.6. Dialect classifier

We trained ten models using different random splits of training and test datasets. Supervised learning was used, with unique labels assigned to each pool of training audios of those birds who inhabit the same region. Then, models were trained to recognize vocal similarities across each geographical population. We calculate the accuracies of models by comparing the predicted classes with the real classes for the test subset. We also evaluate the performance obtained by chance, randomly permuting class labels and computing the corresponding accuracy. We generated ROC curves and a confusion matrix for the pool of predictions. The ROC curve plots the true positive rate (TPR) against the false positive rate (FPR) at different threshold settings, and can illustrate the diagnostic ability of a binary classifier as the threshold is varied. Since our problem is multi-class, we used the one-versus-all strategy to create the ROC curves for each class, defining the class of interest as positive and any other class as negative. The AUC score, the area under the ROC curve, serves as a metric of the model’s performance, with higher values indicating better performance. Additionally, we calculated a confusion matrix, which shows the true labels versus predicted labels, normalized by the total examples for each true class. The diagonal elements of the confusion matrix represent the success rates for each class, while the off-diagonal elements represent the rates of misclassification. Finally, to better evaluate the classification performance of our models, we calculated precision, recall, and F1-score from the predictions of the ten models. Precision is defined as the ratio of true positives of a class to the total number of times that the classifier predicted that class, indicating the level of confidence in the classifier’s ability to correctly identify all members of the positive class. Recall is defined as the ratio of true positives of a class to the total number of actual members of the positive class, indicating the proportion of positive instances that were correctly classified. The F1-score is a harmonic mean of these two indices, providing a balanced measure of classification performance. We compared these performance values for each class (i.e. each region) with those obtained by chance after randomly permuting the labels.

### 2.7. Individual classifier

For individual bird identification, we used the same random splits of training and test data sets as used for the dialect classifier. We trained ten models with supervised learning, from bird identity labels and the pool of training spectrogram chunks of each bird. Then, models were trained to recognize vocal properties of each individual. We compute accuracies of models by comparing the predicted classes with the real classes for the test subset. We also evaluate the performance obtained by chance, randomly permuting class labels and computing the corresponding accuracy. We then joined predictions obtained from models for a robust evaluation of classifier errors and confusion matrix. We compute the average prediction confusion within a region and outside a region as the mean of classification errors made by the predictions of the model in the identification of an individual with another individual that could be from the same region or from different region, respectively. We compared both distributions for each population.

## 3. Results

### 3.1. Dialects

We obtained classifier accuracies of (0.84 +/- 0.04) and compared accuracy values distribution with that obtained by chance (0.30 +/- 0.02). A paired sample t-test showed significant differences between distributions (p-value=2.82e-11), reflecting significantly higher accuracies for the model than those obtained after random permutation of labels (Figure 3.a).

The model showed good performances in prediction for most classes according to ROC curves (Figure 3.b) and confusion matrix (Figure 3.c), with the exception of SL class, for which showed fine performance but with undesirable confusions mostly with Co class. On further inspection, the misclassifications of San Luis chunks belonged mostly to songs of one unique individual, while the other two individuals showed error rates close to those expected based on the average error rate obtained for individuals from all populations. The data of this particular individual consisted in only three (the minimum) songs of not very long durations and spectrograms with not too intensive but extended gray blurs, reflecting not totally removed background noise in different frequency ranges for each song, that could be affecting the prediction performance.

We then analyze the model’s performance by using Precision, Recall, and F1-score. Shapiro-Wilk tests did not reject the null hypothesis that the differences between pairs are normally distributed for most values, but rejected for few distributions. Notwithstanding, both paired sample t-tests and Wilcoxon signed-rank tests reflected significant higher performance values of the model than that obtained by chance for all regions (p-values < 0.05: Figure 3.d).

Performance values of the model presented differences between populations, but also did the performance values obtained by chance. This could be a consequence of the well-known sensibility of these metrics to data imbalance. Hence, we evaluate the relation between performance values and the amount of data used for testing the model (which is similar to that present in the full data since splitting was made in a stratified way). Shapiro-Wilk and Levene tests rejected the null hypothesis of normality and homoscedasticity of distributions. Significant Spearman’s rank correlation was found between performance values and amount of data for Recall (rho=0.55; p-value=2.41e-4) and F1-score (rho=0.52; p-value=5.92e-4), but not for Precision (rho=-0.09; pvalue=0.56). This indicates an effect of data imbalance on model performances. However, results from randomly permuted labels shown significantly higher Spearman’s correlations for Precision (rho=0.90; p-value=2.17e-15), Recall (rho=0.85; p-value=3.10e-12), and F1-score (rho=0.88; p-value=6.34e-14). Thus, differences in performance values between populations could show an effect of data imbalance, but are not fully explained by it.

### 3.2. Individual identification

We obtained classifier accuracies of (0.63 +/-0.03) and compared accuracy values distribution with that obtained by chance (0.06 +/-0.01). A paired sample t-test showed significant differences between distributions (p-value=3.40e-13), reflecting significantly higher accuracies for the model than those obtained after random permutation of labels (Figure 4.a).

**Figure 4:**
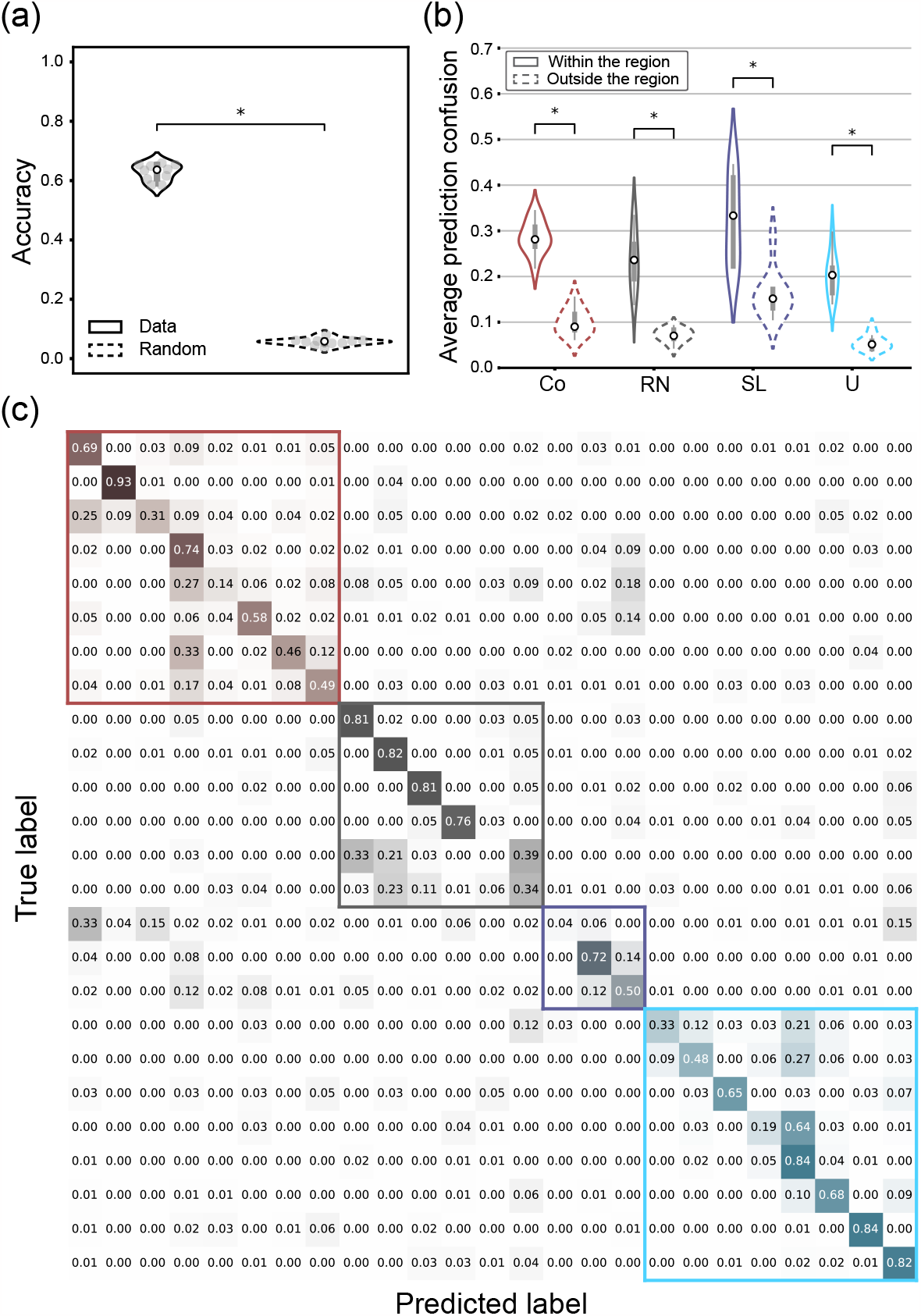
Results of individual classifier. Population color code used for plotting (Co: red; RN: black; SL: blue; U: sky blue) for the regions of Corrientes (Co), San Luis (SL), Rio Negro (RN) and Uruguay (U). (a) Accuracies obtained for test data and random labeled test data, reflecting significant higher values for the model (*; p-value=3.40e-13) by comparing with a paired sample t-test. (b) Average prediction errors grouped in violin plots to differentiate confusions in individuals classification with other individuals within the same region (solid line) or with individuals from other regions (dashed line) for each population. Results shown significantly higher confusions within a population than among populations in all cases, according to Wilcoxon rank sums tests (*; p-value < 0.05), which reflects the presence of dialects. (c) Confusion matrix obtained by the classifier. Diagonal elements of matrix highlight the good performance of the model in identification of individuals for most of the cases but not for all. The non-diagonal elements of the matrix showed that the highest confusions were made between individuals from the same population, as the confusion matrix is closer to a block-diagonal form with blocks separating individuals from each region.

Since a fully random permutation of labels does not prevent the incidence of songs similarities across regions on the individual identification, i.e. individual recognition could be inflated by songs likeness within a dialect, in accordance to results obtained in section 3.1., we performed a random permutation between labels of individuals from the same geographical regions for the evaluation of the model. This constrained random permutation of labels resulted in an accuracy value of (0.15 +/-0.02). We compared it with model accuracy distribution. A paired sample t-test, also showed significant differences between distributions (p-value= 3.08e-12), and accuracies significantly higher for the model.

Next, we analyzed how frequently the model misclassified an individual’s identity and assigned the label of another individual. We compared cases where the incorrect label belonged to an individual within the same population to those where it belonged to an individual in a different population. To do this, we computed the average confusion in predicting an individual’s identity within and outside their region. Shapiro-Wilk tests rejected the null hypothesis of normality of within and among regions average prediction confusions only for SL population. Levene tests rejected the null hypothesis of homoscedasticity for all populations except for Co. Wilcoxon rank sums tests reflected significantly higher values of average prediction errors of the model within regions than among regions for all the populations (p-values < 0.05, Figure 4.b). This reflects that the classifier of individuals is more propense to commit errors in discrimination between individuals that inhabit the same geographical region.

The model showed good performances in individual’s identification for most of the individuals as the confusion matrix, obtained for the pool of predictions, reflects a visible proximity to a diagonal form (Figure 4.c). Notwithstanding, the model showed worse, although not so low, performances for some individuals and a pair of isolated cases for which the capacity in discrimination of individuals was practically null. However, most of the highest confusions in identification of individuals were made between individuals of the same population, which is remarked by the almost block-diagonal form of confusion matrix, where the blocks separate individuals of each region. These results, quantified in Figure 4.b., reinforce the evidence for the presence of dialects in the vocalizations of the Yellow Cardinal obtained in section 3.1.

## 4. Discussion

Models trained using data labeled by regions and individuals had a better-than-chance performance in recognizing dialects and individuals, respectively. This indicates that deep learning classifiers can achieve relatively high accuracy even for an endangered species with limited data availability due to a small number of individuals and songs. Our main finding is that the intra-individual variability in Yellow Cardinal songs is lower than the inter-individual variability, which has not been reported before to our knowledge. This suggests that deep learning classifiers can utilize these differences for individual identification.

The dialects classifier showed good performance for most populations, but it struggled with predicting songs from the San Luis (SL) population, often assigning them the label of the Corrientes (Co) population. Previous research comparing acoustic features of Yellow Cardinal vocalizations between these two populations did not find statistical differences (Domínguez et al., 2016), but a discriminant analysis correctly differentiated them. One possible explanation for the confusion in the classifier could be the low sample size of the San Luis population, although this is not the only factor affecting the model’s performance. It is also possible that the importance for the classification of background noise artifacts, which could not be completely removed with the noise reduction preprocessing, is being amplified by the effect of data imbalance and affecting the performance of the models. Therefore, exploring alternative pre-processing methods could improve the performance of future models.

Although the classifier of individuals was trained for identification of individuals, an additional and interesting result was revealed. The model was more prone to confound predictions between individuals from the same population than between individuals from different populations. This implies that *the model was capable of identifying dialects even when that was not the task for which it was trained*. Hence, deep learning methods applied to audio-based analysis used here, brings out that vocal signatures of Yellow Cardinal reflect a hierarchical structure in the characteristics of vocalizations. This could be analogous for several species of songbirds, for which songs could have a variability that defines different levels of discrimination of vocal characteristics, allowing recognition of species, dialects, or individuals at different scales.

In this work, we trained the classifier of individuals using time and frequency representations of the bird songs, defined by the spectrograms instead of acoustic features (Budka et al., 2015; Xia et al., 2012). This approach has the advantage of allowing more general applications of the method with less manual intervention required and dependence on data. Since the problem is basically a pattern recognition problem in an image, we used data augmentation as used for visual tasks. However, we constrain the augmentation techniques to be compatible with the possible acoustic and sampling properties. Although this method plead for increase training data without loss of biological sense, an outperformer approach could be data augmentation based on bird vocal production models (Gardner et al., 2001; Mindlin, 2017), which have shown to be a good approach for individual discrimination tasks (Bistel et al., 2022a,b; Tubaro and Mindlin, 2019), avoiding noise reduction issues by parameterizing acoustic gestures. This parameterization could allow a method of pitch extraction useful for coping with soundscape noise sources which are hard to be smoothed with our approach, and also could contribute to a better description of the proper biological variability of vocalizations.

The endangered status of Yellow Cardinal (BirdLife International, 2018) in addition with the several factors threatening its preservation, which impose a seemingly trend to persistent population decline (Domínguez et al., 2020; Reales et al., 2019), and the reported significant reduction of its reproductive success (Domínguez et al., 2015), reflect the necessity of stronger and faster strategies of conservation. Noninvasive automatic bioacoustic monitoring of endangered or rare bird species with machine learning models could help investigate spatial temporal distributional shifts at a finer scale. In this way, species responses to habitat modification or climate change could be detected earlier.

Although this requires important efforts in the short term, it will allow more ambitious projects to be tackled with less investment, and to focus on advanced applications with a potentially crucial role in post-research, i.e. concrete policies that are needed. In that regard, our results could be joined with that obtained by genetic studies (Domínguez et al., 2019, 2017), in order to re-evaluate management units that preserve genetic and cultural variation in this endangered species, for further application on suitable and successful reintroduction programmes for this species.

As future work, classifiers could be enhanced by creating synthetic songs through bird vocal production models, considering compatible biological variability of vocalizations. These synthetic songs could also be used for playback experiments to identify the specific parameters that allow individuals to recognize dialects. This could be relevant for better describing cultural units, which are very sensitive to the successful reinsertion of recovered individuals of threatened species, allowing for greater possibilities for their conservation.

## 5. Conclusions

We applied deep learning techniques to audio-based analysis of Yellow Cardinal songs from field recordings. This is the first time that this approach has been applied to this threatened species. We found evidence of vocal signatures of Yellow Cardinal, which allows discrimination between dialects and individuals. The results obtained reflect that these vocal signatures present higher inter-individual variability than intraindividual variability, allowing individual discrimination, but a higher variability between subjects from different populations, allowing dialect discrimination. Our approach not only confirmed previously reported evidence of dialects, but also provides a plausible automated method for characterizing cultural units within the species. This information, combined with genetic data, may aid in the development of strategies for successful reintroduction programs of recovered individuals in threatened species. Additionally, our innovative approach to individual discrimination using deep neural networks shows promise for non-invasive acoustic monitoring strategies, which could have a positive impact on conservation efforts for endangered bird species, particularly the Yellow Cardinal. Our study highlights the potential of advanced machine learning techniques as a valuable tool for avian conservation, particularly for rare bird species with limited available data. These findings suggest that further research in this area could have important implications for conservation biology.

## Author contribution statements

MD, BM, JCR collected the data. HB, MD, GBM designed the study. HB, GBM performed the analysis. HB, MD, BM, JCR, GBM discussed the results, wrote, reviewed and edited manuscript.

## Funding

This work was supported by funding from Agencia Nacional de Promocion Cientifica y Tecnologica (FON-CYT, MINCYT) grant PICT 2017-4681. HB is a post-doctoral fellow from ANCyT.

## Declaration of Competing Interest

The authors declare no conflict of interests.

## Data availability

Data available upon request.

## Acknowledgments

GM thanks the URJC for the hospitality and support during his sabbatical stays. The authors acknowledge a grant PICT 2017-4681 from ANCyT. Alvaro Riccetto shared song recordings from Uruguay.

## Notes

### Competing Interest Statement

The authors have declared no competing interest.

